# Stationary distributions and metastable behaviour for self-regulating proteins with general lifetime distributions

**DOI:** 10.1101/2020.04.25.061101

**Authors:** Candan Çelik, Pavol Bokes, Abhyudai Singh

**Affiliations:** Department of Applied Mathematics and Statistics, Comenius University, Bratislava 84248, Slovakia; Mathematical Institute, Slovak Academy of Sciences, Bratislava 81473, Slovakia; Department of Electrical and Computer Engineering, University of Delaware, Newark, Delaware 19716, USA

**Keywords:** stochastic gene expression, master equation, stationary distribution, metastable systems

## Abstract

Regulatory molecules such as transcription factors are often present at relatively small copy numbers in living cells. The copy number of a particular molecule fluctuates in time due to the random occurrence of production and degradation reactions. Here we consider a stochastic model for a self-regulating transcription factor whose lifespan (or time till degradation) follows a general distribution modelled as per a multidimensional phase-type process. We show that at steady state the protein copy-number distribution is the same as in a one-dimensional model with exponentially distributed lifetimes. This invariance result holds only if molecules are produced one at a time: we provide explicit counterexamples in the bursty production regime. Additionally, we consider the case of a bistable genetic switch constituted by a positively autoregulating transcription factor. The switch alternately resides in states of up- and downregulation and generates bimodal protein distributions. In the context of our invariance result, we investigate how the choice of lifetime distribution affects the rates of metastable transitions between the two modes of the distribution. The phase-type model, being non-linear and multi-dimensional whilst possessing an explicit stationary distribution, provides a valuable test example for exploring dynamics in complex biological systems.

## 1 Introduction

Biochemical processes at the single-cell level involve molecules such as transcription factors that are present at low copy numbers [6, 46]. The dynamics of these processes is therefore well described by stochastic Markov processes in continuous time with discrete state space [15, 22, 42]. While few-component or linear-kinetics systems [16] allow for exact analysis, in more complex system one often uses approximative methods [12], such as moment closure [4], linear-noise approximation [3, 9], hybrid formulations [25, 26, 33], and multi-scale techniques [38, 39].

In simplest Markovian formulations, the lifetime of a regulatory molecule is memoryless, i.e. exponentially distributed [10, 47]. However, transcription factors are complex macromolecules, which can be present in various molecular conformations, and whose removal can require a complex interplay of multiple pathways [13, 37]. Therefore, their lifetime distributions can assume far more complex forms than the simple exponential.

In Section 2, we formulate, both in the deterministic and stochastic settings, a one-dimensional model for the abundance of a transcription factor with a memoryless lifetime. Since many transcription factors regulate their own gene expression [2], we allow the production rate to vary with the copy number. We show that the deterministic solutions tend to the fixed points of the feedback response function; in the stochastic framework, we provide the stationary distribution of the protein copy number.

In Section 3, we proceed to characterise the steady-state behaviour of a structured model that accounts for complex lifetime pathways. The model is multidimensional, each dimension corresponding to a different class and stage of a molecule’s lifetime; the chosen structure accounts for a wide class of phasetype lifetime distributions [34, 45]. We demonstrate that the deterministic fixed points and the stochastic stationary distribution that were found for the onedimensional framework remain valid for the total protein amount in the multidimensional setting.

We emphasise that the distribution invariance result rests on the assumption of non-bursty production of protein. The case of bursty production is briefly discussed in Section 4, where explicit counter-examples are constructed by means of referring to explicit mean and variance formulae available from literature for systems without feedback [28, 36].

In the final Section 5, we approximate the stochastic protein distribution by a mixture of Gaussians with means at deterministic fixed points and variances given by the linear-noise approximation [8, 30]. Additionally, we study the rates of metastable transitions [40, 43] between the Gaussian modes in the one-dimensional and structured settings.

## 2 One-dimensional model

### Deterministic framework

The dynamics of the abundance of protein *X* at time *t* can be modelled deterministically by an ordinary differential equation

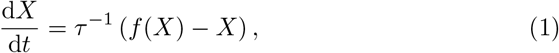

which states that the rate of change in *X* is equal to the difference of production and decay rates. The decay rate is proportional to *X*; the factor of proportion-ality is the reciprocal of the expected lifetime *τ*. The rate of production per unit protein lifetime is denoted by *f* (*X*) in (1); the dependence of the production rate on the protein amount *X* implements the feedback in the model. Equating the right-hand side of (1) to zero yields

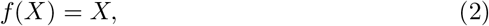

meaning that steady states of (1) are given by the fixed point of the production response function *f* (*X*).

### Stochastic framework

The stochastic counterpart of (1) is the Markov process with discrete states 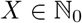 in continuous time with transitions *X* → *X* — 1 or *X* → *X* +1, occurring with rates *X*/*τ* and *f* (*X*)/*τ* respectively (see the schematic in Figure 1). Note that in case of a constant production rate, i.e. *f* (*X*) ≡ λ, the model turns into the immigration-and-death process [32]; in queueing theory this is also known as *M*/*M*/∞ queue [21]. The stationary distribution of the immigration–death process is known to be Poissonian with mean equal to λ [32].

**Figure 1:**
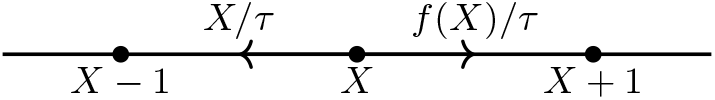
A diagram of the one-dimensional model. The number of molecules *X* can decrease by one or increase by one. The stochastic rates (or propensities) of these transitions are indicated above the transition edges.

For a system with feedback, the probability *P* (*X*, *t*) of having *X* molecules at time *t* satisfies the master equation

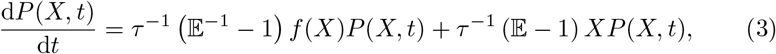

in which 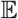 is the van-Kampen step operator [30]. Inserting *P*(*X,t*) = *π*(*X*) into (3) and solving the resulting difference equation, one finds a steady-state distribution in the explicit form

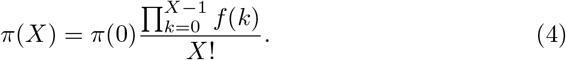

The probability *π*(0) of having zero molecules plays the role of the normalisation constant in (4), which can be uniquely determined by imposing the normalisation condition *π*(0) + *π*(1) +… = 1. Note that inserting *f* (*X*) ≡ λ into (4) results in the aforementioned Poissonian distribution with *π*(0) = e^−λ^.

## 3 Multiclass–multistage model

In this section, we introduce a structured multiclass–multistage model which is an extension of one-dimensional model introduced in the previous section. The fundamentals of the multidimensional model are as shown in Figure 2. A newly produced molecule is assigned into one of *K* distinct classes. Which class is selected is chosen randomly according to a discrete distribution *p*_1_,…, *p_K_*, The lifetime of a molecule in the *i*-th class consists of *S_i_* stages. The holding time in any of these stages is memoryless (exponential), and parametrised by its mean *τ_ij_*, where *i* indicates which class and *j* indicates which stage. Note that

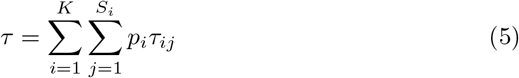

gives the expected lifetime of a newly produced molecule. After the last (*S_i_*-th) stage, the molecule is degraded. The total distribution of a molecule lifetime is a mixture, with weights *p_i_*, of the lifetime distributions of the individual classes, each of which is a convolution of exponential distributions of the durations of the individual stages; such distributions are referred to as phase-type distribution and provide a wide family of distribution to approximate practically any distribution of a positive random variable [45].

**Figure 2:**
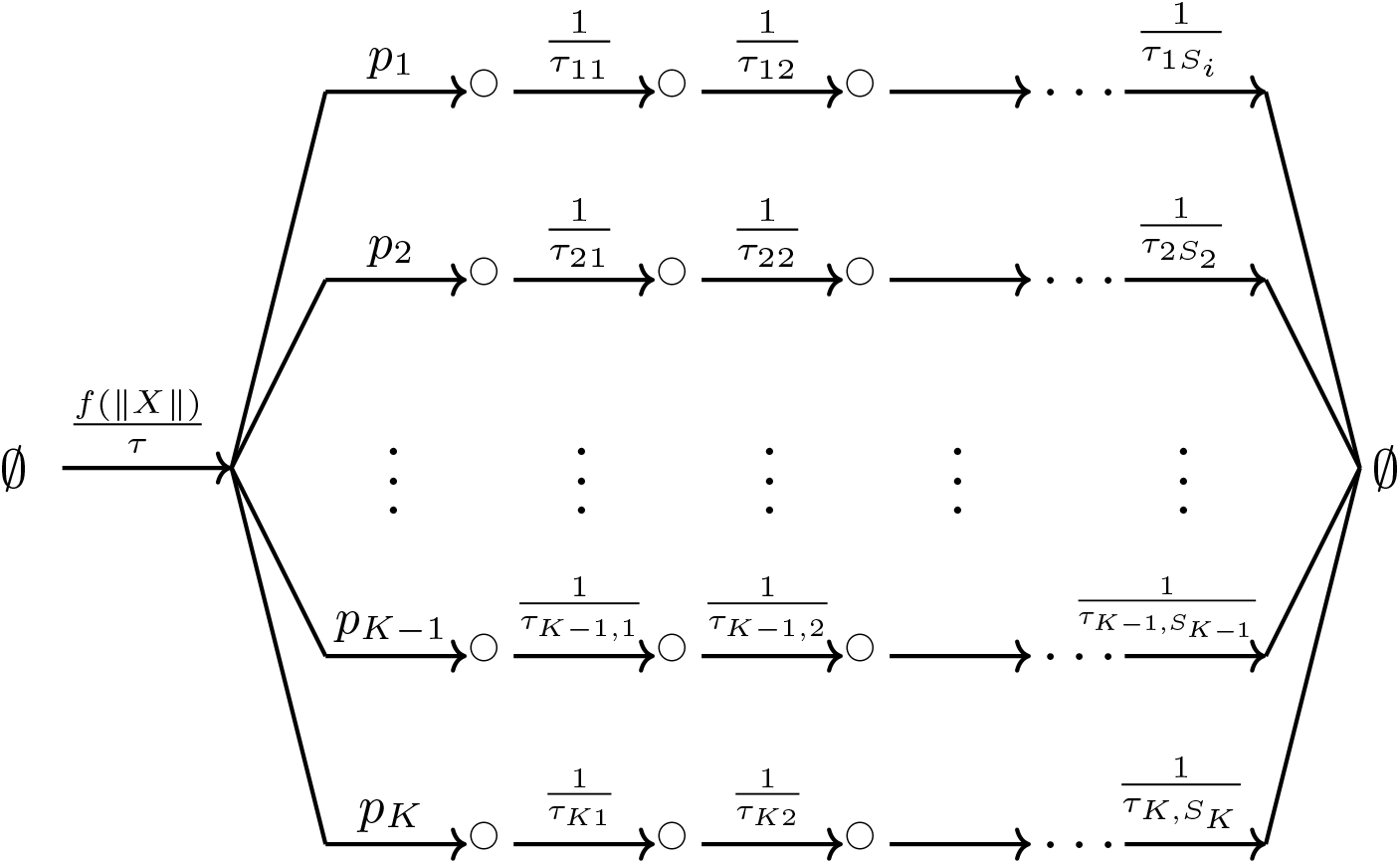
A schematic representation of multiclass–multistage model. A newly produced molecule is randomly assigned, according to a prescribed distribution *p_1_*,…,*p_κ_*, into one of *K* distinct classes. The lifetime of a molecule in the *i*-th class consists of *S_i_* consecutive memoryless stages, and ends in the degradation of the molecule. The expected holding time in the j-th stage of the *i*-th class is *τ_ij_*. The production rate is a function of the total number ǁ*X*ǁ of molecules across all stages and classes.

We denote by *X_ij_* the number of molecules in the *i*-th class and the *j*-th stage of their lifetime, by

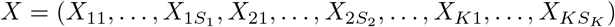

the 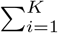 *S_i_*-dimensional copy-number vector, and by

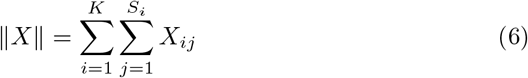

the total number of molecules across all classes and stages.

### Deterministic framework

The deterministic description of the structured model is given by a system of coupled ordinary differential equations

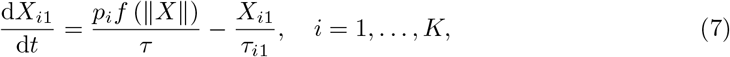

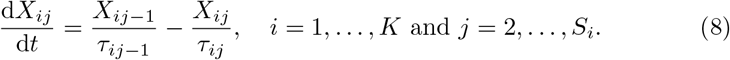

The right-hand sides of (7)–(8) are each equal to the difference of appropriate arrival and departure rates at/from a particular compartment of the structured model. The departure rates are proportional to the number of molecules in the compartment, with the reciprocal of the holding time giving the factor of proportionality. The arrival rate takes a different form for the first stages (7) and for the other stages (8). For the first stage, the arrival is obtained by the product of the production rate *f* (ǁ*X*ǁ)/*τ* and the probability *p_i_* of selecting the *i*-th class. For the latter stages, the arrival rate is equal to the departure rate of the previous stage.

Equating (7)–(8) to zero, we find that

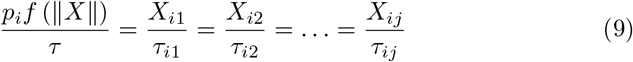

hold at steady state, from which it follows that

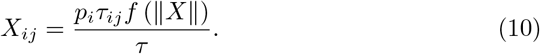

Summing (10) over *i* = 1,…, *K* and *j* = 2,…, *S_i_*, and using (5) and (6), yield

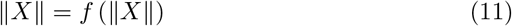

for the total protein amount (6). Thus, the protein amount at steady state is obtained, like in the one-dimensional model, by calculating the fixed points of the feedback response function.

Combining (11) and (9) we find

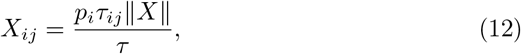

which means that at steady state the total protein amount is distributed among the compartments proportionally to the product of class assignment probability and the mean holding time of the particular compartment.

### Stochastic framework

Having shown that the stationary behaviour of the onedimensional and the structured multi-dimensional models are the same in the deterministic framework, we next aim to demonstrate that the same is also true in the stochastic context. Prior to turning our attention to the feedback system, it is again instructive to discuss the case without regulation, i.e. *f*(ǁ*X*ǁ) ≡ λ; the new molecule arrivals are then exponentially distributed. In the language of queueing theory, the process can be reinterpreted as the *M*/*G*/∞ queue with exponential arrivals of customers, a general phase-type distribution of service times, and an infinite number of servers. It is well known that the steady-state distribution of an *M*/*G*/∞ queue is Poisson with mean equal to λ [41]. Thus, without feedback, we obtain the very same Poisson(λ) distribution that applies in the one-dimensional case.

In the feedback case, the probability *P*(*X, t*) of having *X* = (*X*_11_,…, *X_K,S_K__*) copy numbers in the individual compartments at any time *t* satisfies the master equation

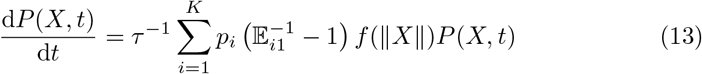

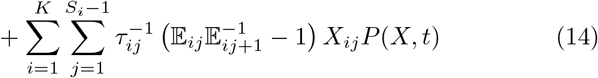

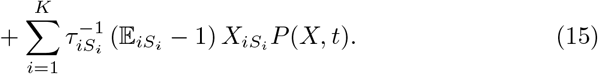

The right-hand-side terms (13), (14), and (15) stand for the change in probability mass function due to the production, moving to next stage, and decay reactions, respectively. Note that 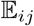 is a step operator which increases the copy number of molecules in the *i*-th class at the *j*-th stage by one [30]. Likewise, 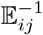 decreases the same copy number by one. Rearrangement of terms in the master equation yields

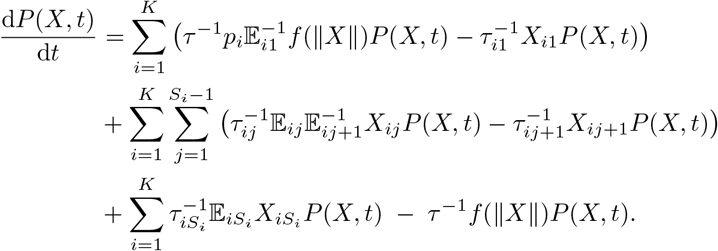

Equating the derivative to zero, we derive for the stationary distribution *π*(*X*) an algebraic system

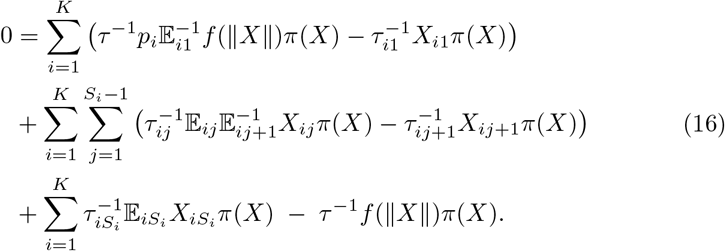

Clearly, it is sufficient that

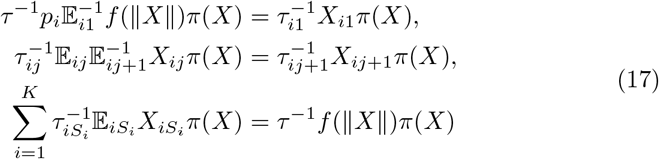

hold for *π*(*X*) in order that (16) be satisfied. One checks by direct substitution that

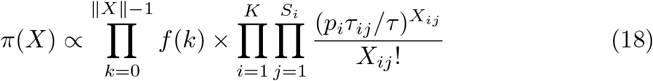

satisfies (17); therefore, (18) represents the stationary distribution of the structured model. In order to interpret (18), we condition the joint distribution on the total protein copy number, writing

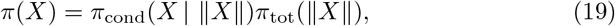

in which the conditional distribution is recognised as the multinomial [29]

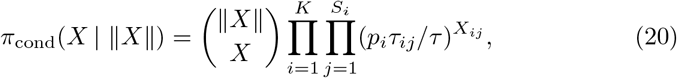

and the total copy number distribution is given by

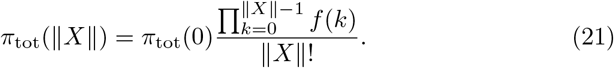

By (20), the conditional means of *X_ij_* coincide with the deterministic partitioning of the total copy number (12). Importantly, comparing (21) to (4), we conclude that the one-dimensional and multi-dimensional models generate the same (total) copy number distributions.

## 4 Bursting

The independence of stationary distribution on the lifetime distribution relies on the assumption of non-bursty production of protein that has implicitly been made in our model. In this section, we allow for the synthesis of protein in bursts of multiple molecules at a single time [14, 17]. Referring to previously published results [28, 36], we provide an counterexample that demonstrates that in the bursty case different protein lifetime distributions can lead to different stationary copy-number distributions. The counterexample can be found even in the absence of feedback.

Let *G*(*t*) = Prob[*T* > *t*] denote the survival function of an individual molecule and let

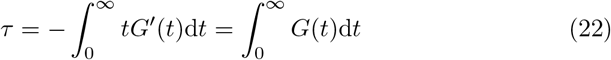

be the mean lifetime. Bursty production means that the number of molecules can change within an infinitesimally small time interval of length d*t* from *X* to *X* + *j*, where *j* ≥ 1, with probability *λτ*^−1^*b_j_* d*t*, in which λ is the burst frequency and *b_j_* = Prob[B = *j*] is the probability mass function of the random burst size *B* variable.

In queueing theory, bursty increases in the state variable are referred to as batch customer arrivals. Specifically, a bursty gene-expression model without feedback and with general lifetime distribution corresponds to the *M^X^*/*G*/∞ queue with memoryless (exponential) batch arrivals, general service distribution, and an infinite number of servers.

Previous analyses [28, 36] show that the steady-state protein mean 〈*X*〉 and the Fano factor *F* = Var(*X*)/〈*X*〉 are given by

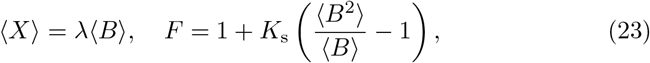

where

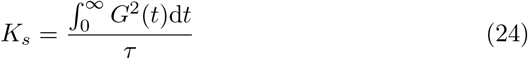

is referred to as the senescence factor. Elementary calculation shows that *K_s_* = 1/2 if the lifetime distribution is exponential with survival function *G*(*t*) = *e^−t/τ^* and that *K_s_* = 1 if the lifetime distribution is deterministic with survival function *G*(*t*) = 1 for *t* < *τ* and *G*(*t*) = 0 for *t* ≥ *τ*. Thus, although two lifetime distributions result in the same value of the stationary mean protein copy number, they give a different value of the noise (the Fano factor); therefore the copy-number distributions are different.

## 5 Metastable transitioning

Transcription factors that self-sustain their gene expression by means of a positive feedback loop can act as a simple genetic switch [5, 20]. A positive-feedback switch can be in two states, one in which the gene is fully activated through its feedback loop, while in the other the gene is expressed at a basal level. The switch serves as a basic memory unit, retaining the information on its initial state on long timescales, and very slowly relaxing towards an equilibrium distribution. It is therefore important to investigate not only the stationary, but also transient distributions, which are generated by a positively autoregulating transcription factor.

Following previous studies [7, 11, 19], we model positive feedback by the Hill function response curve

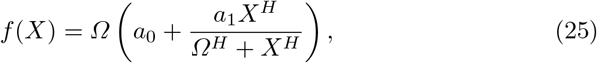

in which *a*_0_ and *a*_1_ represent the basal and regulable production rates, *H* is the cooperativity coefficient, and *Ω* gives the critical amount of protein required for half-stimulation of feedback. Provided that *H* > 1, one can find *a*_0_ and *a*_1_ such that (25) possesses three distinct fixed points *X*_−_ < *X*_0_ < *X*_+_, of which the central is unstable and the other two are stable (Figure 3, left). The two stable fixed points provide alternative large-time outcomes of the deterministic models (1) and (7)–(8).

**Fig. 3:**
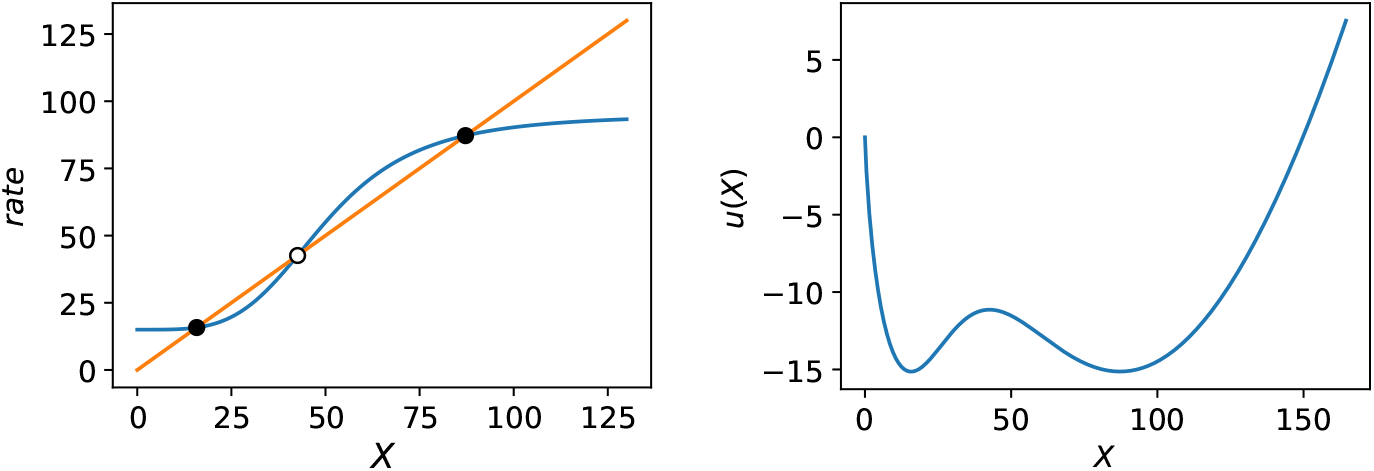
*Left:* A sigmoid feedback response function (blue curve) intersects the diagonal (orange line) in multiple fixed points. Ones that are stable to the rate equation (1) (full circles) are interspersed by unstable ones (empty circle). *Right:* The potential *u*(*x*), defined by (33), is a Lyapunov function of the rate equation (1). The local minima, or the troughs/wells, of the potential are situated at its stable fixed points; the local maximum, or the barrier, of the potential coincides with the unstable fixed point. *Parameter values for both panels:* We use the Hill-type response (25) with = 0.3, *a*_1_ = 1.6, *H* = 4, *Ω* = 50.

Bistability of deterministic models translates into bimodal distributions in the stochastic framework. For large values of *Ω*, the bimodal protein distribution can be approximated by a mixture of Gaussian modes which are located at the stable fixed points *X*_±_ (see Figure 4),

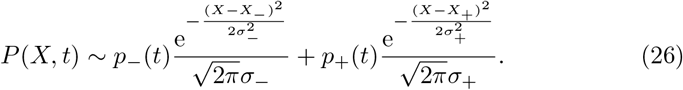

**Fig. 4:**
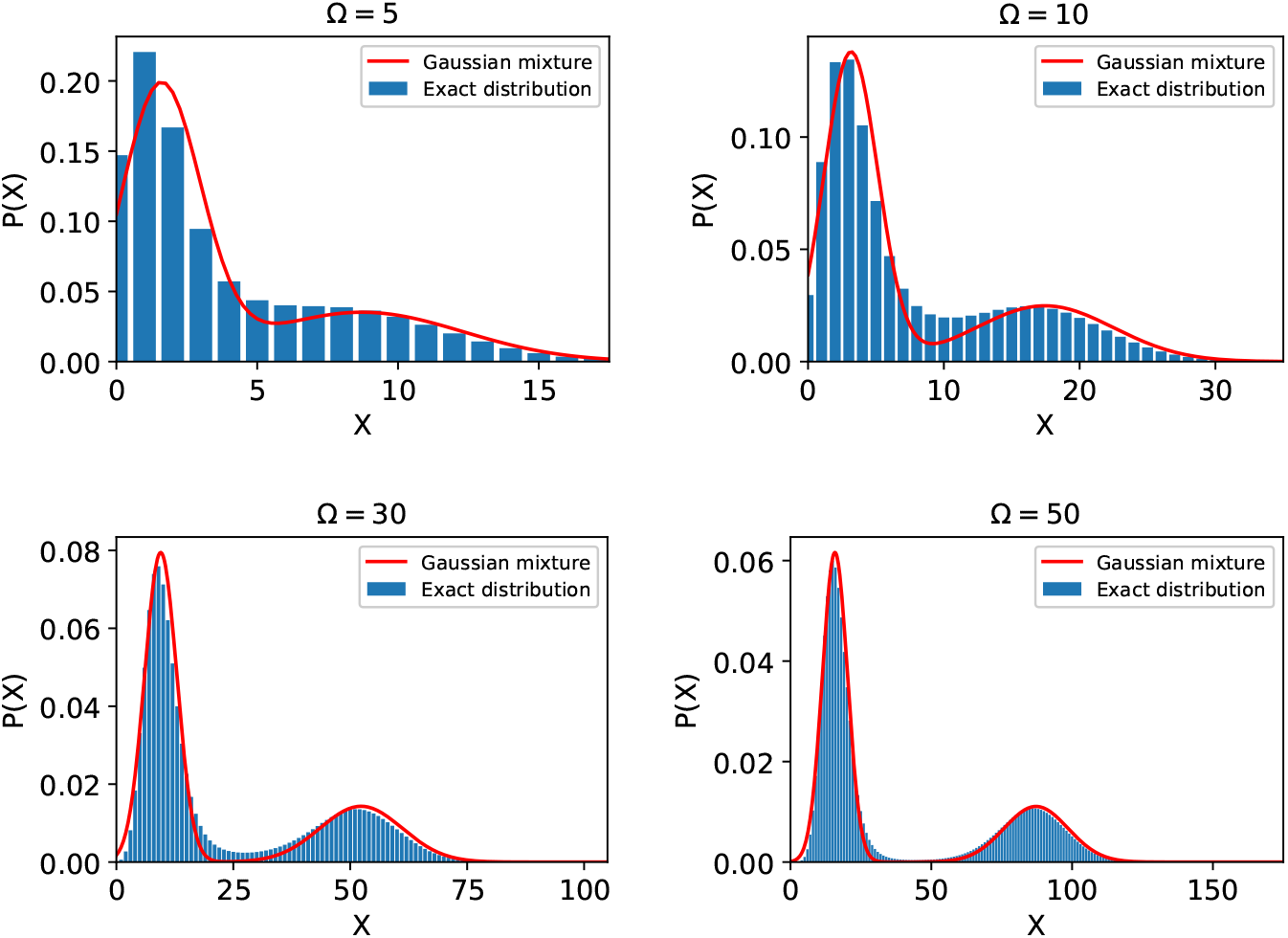
Exact stationary protein distribution (4) and the Gaussian-mixture approximation (26) in varying system-size conditions. The means of the Gaussians are given by the stable fixed points of *f*(*X*); the variances are given by linear-noise approximation (27). The mixture weights are given by *p*_+_(∞) = *T*_+_/(*T*_+_ + *T*_−_), *p*_−_(∞) = *T*_−_/(*T*_+_ + *T*_−_), where the residence times are given by the Arrhenius-type formula (32). We use a Hill-type response (25) with *a*_0_ = 0.3, *a*_1_ = 1.6, *H* = 4, and *Ω* shown in panel captions.

The mixture approximation (26) is determined not only by the locations *X*±, but also on the variances 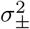 and the weights *p*±(*t*) of the two modes (which are given below). The weights in (26) are allowed to vary with time in order to account for the slow, metastable transitions that occur between the distribution modes.

The invariance result for stationary distributions derived in the preceding sections implies that, in the limit of *t* → ∞, the protein distribution (26) becomes independent of the choice of the protein lifetime distribution. In particular, the same variances 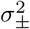 and the same limit values *p*±(∞) of the weights will apply for exponentially distributed and phase-type decay processes. In what follows, we first consult literature to provide results 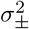 and *p*±(*t*) that apply for the one-dimensional model with exponential decay. Next, we use stochastic simulation to investigate the effect of phase-type lifespan distributions on metastable dynamics.

The variances of the modes are obtained by the linear-noise approximation [35, 44] of the master equation (3), which yields

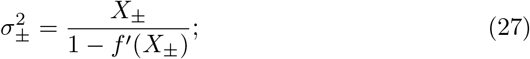

the right-hand side of (27) is equal to the ratio of a fluctuation term (equal to the number of molecules) to a dissipation term (obtained by linearising the rate equation (1) around a stable fixed point).

The metastable transitions between the distribution modes can be described by a random telegraph process (cf. Figure 5, left)

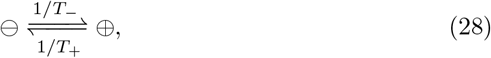

in which the lumped states ⊖ and ⊕ correspond to the basins of attractions of the two stable fixed points; *T*_−_ and *T*_+_ are the respective residence times. The mixture weights *p*_−_(*t*) and *p*_+_(*t*) in (26) are identified with the probabilities of the lumped states in (28); these satisfy the Chapman–Kolmogorov equations

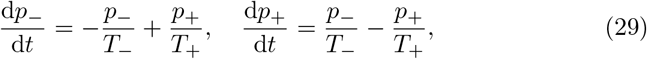

which admit an explicit solution

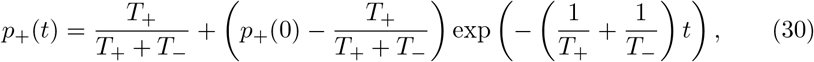

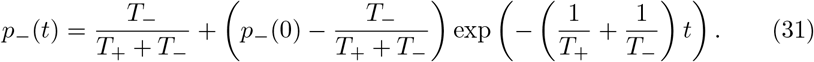

**Figure 5:**
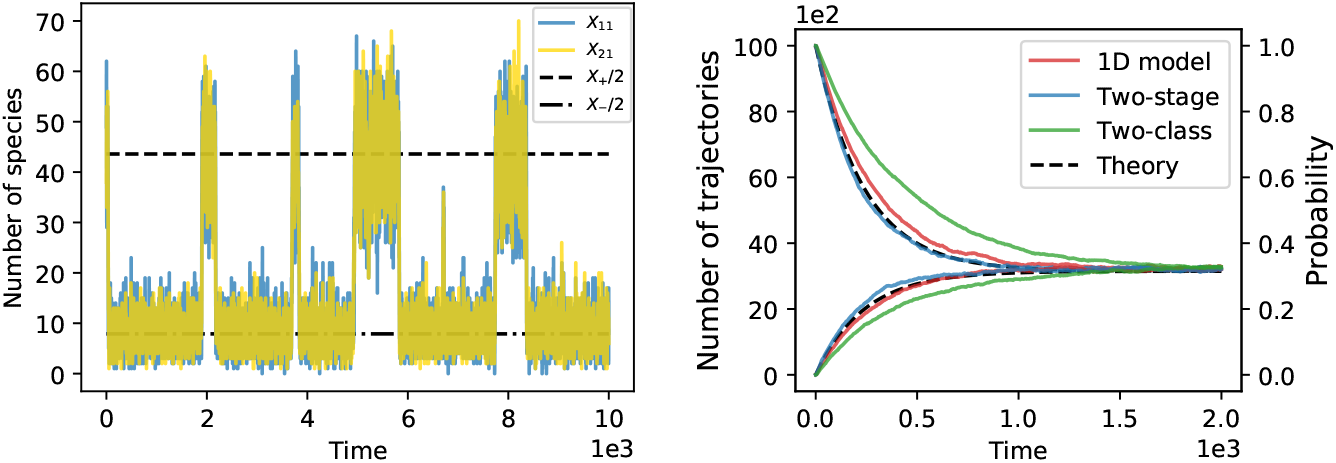
*Left:* Large-time stochastic trajectories of a structured two-class model with parameters as given below. The horizontal lines represent deterministic fixed points as given by (11)−(12). *Right:* The number of trajectories, out of 10^4^ simulation repeats, that reside in the basin of attraction of the upper stable fixed point as function of time. Simulation is initiated at the upper stable fixed point (the decreasing function) or at the lower stable fixed point (the increasing function). The dashed black curve gives the theoretical probability (30) with initial condition *p*_+_(0) = 1 (the decreasing solution) or *p*_+_(0) = 0 (the increasing solution). *Parameter values:* The Hill-function parameters are: *Ω* = 50, *H* = 4, *a*_0_ = 0.3, *a*_1_ = 1.6. The mean lifetime is *τ* = 1. The two-stage model parameters are: *K* = 1, *p*_1_ = 1, *S*_1_ = 2, *τ*_11_ = *τ*_12_ = 0.5. The two-class model parameters are: *K* = 2, *p*_1_ = 1/6, *p*_2_ = 5/6, *S*_1_ = *S*_2_ = 1, *τ*_11_ = 3, *τ*_21_ = 3/5.

The initial probability *p*_+_(0) = 1 — *p*_−_(0) is set to one or zero in (30)−(31) depending on whether the model is initialised in the neighbourhood of the upper or the lower stable fixed point.

With (30)−(31) at hand, the problem of determining the mixture weights in (26) is reduced to that of determining the residence times *T*_±_. Previous large-deviation and WKB analyses of the one-dimensional model [18, 23, 24] provide an Arrhenius-type formula

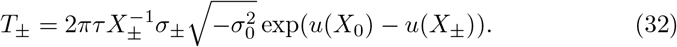

Formula (32) features, on top of the familiar symbols (the mean lifetime *τ*, fixed points *X*_±_ and *X*_0_, linearised variances *σ*_±_, and the Ludolph-van-Ceulen constant *π*), two new symbols: a value 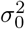 and a function *u*(*X*). The value 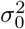 is readily calculated by inserting 0 instead of ± into the fluctuation-dissipation relation (27); note that for the unstable fixed point *X*_0_, the denominator in (27) is negative (cf. Figure 3, left), which renders the whole fraction also negative.

In analogy with the Arrhenius law, the function *u*(*X*) represents an “energy” of state *X*, and is given here explicitly by an indefinite integral [18, 23, 24]

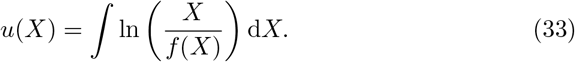

Note that the derivative of (33),

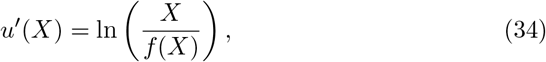

is zero if *f* (*X*) = *X*, i.e. at the fixed points of the feedback response function, is negative if *f* (*X*) > *X* and positive if *f* (*X*) < *X*. Substituting into (33) the solution *X* = *X*(*t*) to the deterministic rate equation (1) and evaluating the time derivative, we find

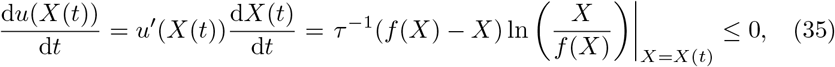

with equality in (35) holding if and only if *X* is a fixed point of the feedback response function *f*(*X*). Therefore, the energy function *u*(*X*) is a Lyapunov function of the ordinary differential equation (1) (Figure 3, right). The exponentiation in (32) dramatically amplifies the potential difference between the stable and the unstable fixed points. For example, a moderately large potential barrier, say 5 (which is about the height of the potential barrier in Figure 3, right), introduces a large factor e^5^ ≈ 150 in (32). This confirms an intuition that metastable transitions between the distribution modes are very (exponentially) slow.

The random telegraph solution (30) is compared in Figure 5 to the residence of stochastically generated trajectories in the basin of attraction of the upper fixed point. The agreement is close for simulations of the one-dimensional model (with an exponential lifetime) and for a structured model with one class and two stages (with an Erlangian lifetime). For a two-class model (with an exponential mixture lifetime), the transitioning also occurs on the exponentially slow timescale, but is perceptibly slower. Sample trajectories were generated in Python’s package for stochastic simulation of biochemical systems GillesPy2 [1]. The one-dimensional model was initiated with ⌊*X*_+_⌋ molecules. The two species in the two-stage and two-class models were initiated to *S* and ⌊*X*_+_⌋ –*S*, where *S* was drawn from the binomial distribution Binom(⌊*X*_+_⌋, 0.5).

## 6 Discussion

In this paper we studied a stochastic chemical reaction system for a self-regulating protein molecule with exponential and phase-type lifetimes. We demonstrated that the exponential and phase-type models support the same stationary distribution of the protein copy number. While stationary distributions of similar forms have previously been formulated in the context of queueing theory [21, 27, 31], our paper provides a self-contained and concise treatment of the onedimensional model and the multi-dimensional structured model that is specifically tailored for applications in systems biology.

We showed that the invariance result rests on the assumption of non-bursty production of protein. We demonstrated that, in the presence of bursts, exponential and deterministic lifetimes generate stationary protein-level distributions with different variances.

Deterministic modelling approaches are used in systems biology as widely as stochastic ones. Therefore, we complemented the stationary analysis of the stochastic Markov-chain models by a fixed-point analysis of deterministic models based on differential equations. The result is that, irrespective of lifetime distribution, the deterministic protein level is attracted, for large times, to the stable fixed points of the feedback response function. Connecting the stochastic and deterministic frameworks, we demonstrated that the stationary distribution of the Markovian model is sharply peaked around the fixed points of the deterministic equation. We showed that the distribution can be approximated by a mixture of Gaussian modes with means given by the deterministic fixed points and variances that are consistent with the traditional linear-noise analysis results.

Next, we focused on the transitions between the distribution modes. These occur rarely with rates that are exponentially small. We compared an asymptotic result, derived in previous literature for the one-dimensional model, to stochastic simulation results of the one-dimensional model and two specific structured models: we chose a model with one class and two stages and a model with two classes each with one stage. The simulation results of the one-dimensional and two-stage models agreed closely to the theoretical prediction; intriguingly, the agreement with theory was closer for the two-stage model. On the other hand, a two-class model showed slower transitioning rates. The theoretical asymptotic results have been derived in [18, 23, 24] only for the one-dimensional model. Large deviations in multi-dimensional models are much harder to quantify that one-variable ones. We believe that the current model, being multi-dimensional while possessing a tractable steady-state distribution, provides a convenient framework on which such methodologies can be developed.

In summary, our study provides an invariance-on-lifetime-distribution result in the deterministic and stochastic contexts for a non-bursty regulatory protein. While the main results concern the stationary behaviour, our study also performs simulation, and opens avenue for future enquiries, into the transient transitioning dynamics.

